# Assessment of the aging of the brown adipose tissue by ^18^F-FDG PET/CT imaging in the progeria mouse model Lmna^G609G/G609G^

**DOI:** 10.1101/220061

**Authors:** Zhengjie Wang, Xiaolong Xu, Yi Liu, Yongheng Gao, Ennui Ma, Welding Yang, Fei Kang, Baohua Liu, Jing Wang

## Abstract

Brown adipose tissue (BAT) is an important energy metabolic organ that is closely related to obesity, type 2 diabetes, and atherosclerosis. Aging is one of the most important determinants of BAT activity. In this study, we used ^18^F-FDG PET/CT imaging to assess the aging of the BAT in Lmna^G609G/G609G^ mice. To evaluate the BAT activity, Lmna^G609G/G609G^ and wild-type (WT) mice were injected with 18F-FDG, and PET/CT imaging was performed. The maximum standardized uptake value (SUV_Max_) of the BAT was measured and the target/nontarget (T/NT) values of BAT were calculated. The transcription and the protein expression levels of the uncoupling protein 1 (UCP1), beta3-adrenergic receptor (β3-AR), and the PRdomain-containing16 (PRDM16), were measured by quantitative real-time polymerase chain reaction (RT-PCR) and Western blotting or immunohistochemical analysis. Apoptosis and cell senescence of the BAT, in WT and Lmna^G609G/G609G^ mice, was detected by Terminal deoxynucleotidyl transferase dUTP nick end labeling (TUNEL), and by CDKN2A/p16INK4a immunohistochemical staining, respectively. At 14 weeks of age, the BAT SUV_Max_ and the expression levels of UCP1, β3-AR and PRDM16 in Lmna^G609G/G609G^ mice was significantly lower than that in WT mice. At the same time, the number of p16INK4a and TUNEL positively stained cells (%) increased in Lmna^G609G/G609G^ mice. Lmna^G609G/G609G^ mice are an ideal model for studying BAT aging. The aging characteristics and the aging mechanism of BAT in Lmna^G609G/G609G^ mice can mimic normal BAT aging.

## Introduction

Aging has been defined as the age-related deterioration of physiological functions of an organism. The essence of aging is the process of gradual decline of the functions of the organ system. Brown adipose tissue (BAT) is an adipose organ which is maintaining core temperature in small mammals and in newborn humans (Cannon and Nedergaard 2004). The functional status of BAT is closely related to obesity, type 2 diabetes and atherosclerosis (Berbee et al. 2015; Koksharova et al. 2017). BAT quality and activity are gradually decrease with age (Pfannenberg et al. 2010).

Aging is one of the most important determinants of BAT activity (Lecoultre and Ravussin 2011). Therefore, it is important to study the relationship between BAT functional status and aging. Since 1996, the researchers conducted long-term study of the relationship between BAT function and aging by measuring BAT-related indicators, including cold-induced heat generation, cold tolerance, body temperature and other macro physiological parameters and in the level of microscopic molecular biomarker such as Uncoupling protein 1 (UCP1), PPAR-g coactivator 1a (PGC1a), PRdomain-containing16 (PRDM16) and beta3-adrenergic receptor (beta3-AR) (Kirov et al. 1996; Lin et al. 2016; Yamashita et al. 1999). However, there are still some problems that cannot be ignored in the above research. Firstly, the research model, as the above study is the normal mouse aging process, and the mouse life of up to 32 weeks, so the research cycle is long, the degree of aging is difficult to unified control(Graber et al. 2015); secondly, the research method, the macroscopic physiological indicators or microscopic molecular markers used in the above studies are indirect data on the metabolic activity of BAT, and often do not accurately reflect the functional status of BAT. Thus, a visualization of the method is necessary to detect BAT metabolic activity of the models of aging.

Since 2003, the presence of adult BAT was confirmed during ^18^F-FDG PET scanning (Cohade et al. 2003). ^18^F-FDG PET/CT is considered to be the “gold standard” of the current BAT function measurement, and the functional status of BAT can be measured directly in vivo by showing glucose metabolism activity (Cypess et al. 2014), and has been widely used in the basics and clinical studies of BAT (Cohade et al. 2003; Yeung et al. 2003). Hutchinson-Gilford progeria syndrome(HGPS) is caused by a mutation in the Lmna gene (De Sandre-Giovannoli et al. 2003a). According to this phenomenon, the HGPS mouse model (Lmna^G609G/G609G^ mice) was constructed and as the models of aging have been widely used in aging studies (Chen et al. 2012; Zhang et al. 2014). However, like any premature aging syndrome, Lmna^G609G/G609G^ is only a partial representation of the multifactorial process of normal aging. The aging of some key organs will not be appearing in Lmna^G609G/G609G^ models such as nervous system (Jung et al. 2012). Whether the BAT function of the Lmna^G609G/G609G^ mice is premature senility; whether the level of ^18^F-FDG uptake is related to the levels of the molecular markers which are associated with the aging of BAT. The above questions directly determine whether the model is suitable for BAT-related aging studies.

In this study, ^18^F-FDG PET/CT imaging technique was used to qualitatively and quantitatively analyze the relationship between ^18^F-FDG PET/CT imaging and aging of BAT in Lmna^G609G/G609G^ mice. The relationship between BAT-related molecular markers UCP1 and β3-AR levels and ^18^F-FDG uptake was examined. In addition, the mechanism of BAT dysfunction in mice was studied.

## Result

### Effects of age on the metabolic activity of BAT in mice

To study the effect of age on Lmna^G609G/G609G^ mice and WT mice, PET/CT imaging was performed from 4 weeks on Lmna^G609G/G609G^ mice and WT mice, once every two weeks and for 16 weeks. As shown in figure 1A, from 4 weeks to 12 weeks, there was no significant difference in SUV_Max_ of BAT between Lmna^G609G/G609G^ mice and WT mice, indicating that FDG uptake in BAT of Lmna^G609G/G609G^ mice was no different from that of WT mice at first 12 weeks. As shown in figure 1B, at 14 weeks of age, ^18^F-FDG uptake of BAT in Lmna^G609G/G609G^ mice was significantly lower than that in WT mice. Statistically, the SUV_Max_ value and the T/NT ratio of BAT did not differ significantly between Lmna^G609G/G609G^ mice and WT mice at 4 weeks of age (SUV_Max_ value: 9.260±0.571 vs. 9.373±0.709, p=0.9069; T/NT ratio:32.27±2.278 vs. 30.41±2.411, p=0.6223) and at 14 weeks of age, the Lmna^G609G/G609G^ mice were significantly lower than WT mice (SUV_Max_ value: 3.673±0.4613 vs.10.34±0.8663, p=0.0003 T/NT ratio: 11.61±1.975 vs. 33.16±2.687, p=0.0030). These findings confirm that the BAT ^18^F-FDG uptake in Lmna^G609G/G609G^ mice decreased significantly at 14 weeks of age.

**Figure 1.**
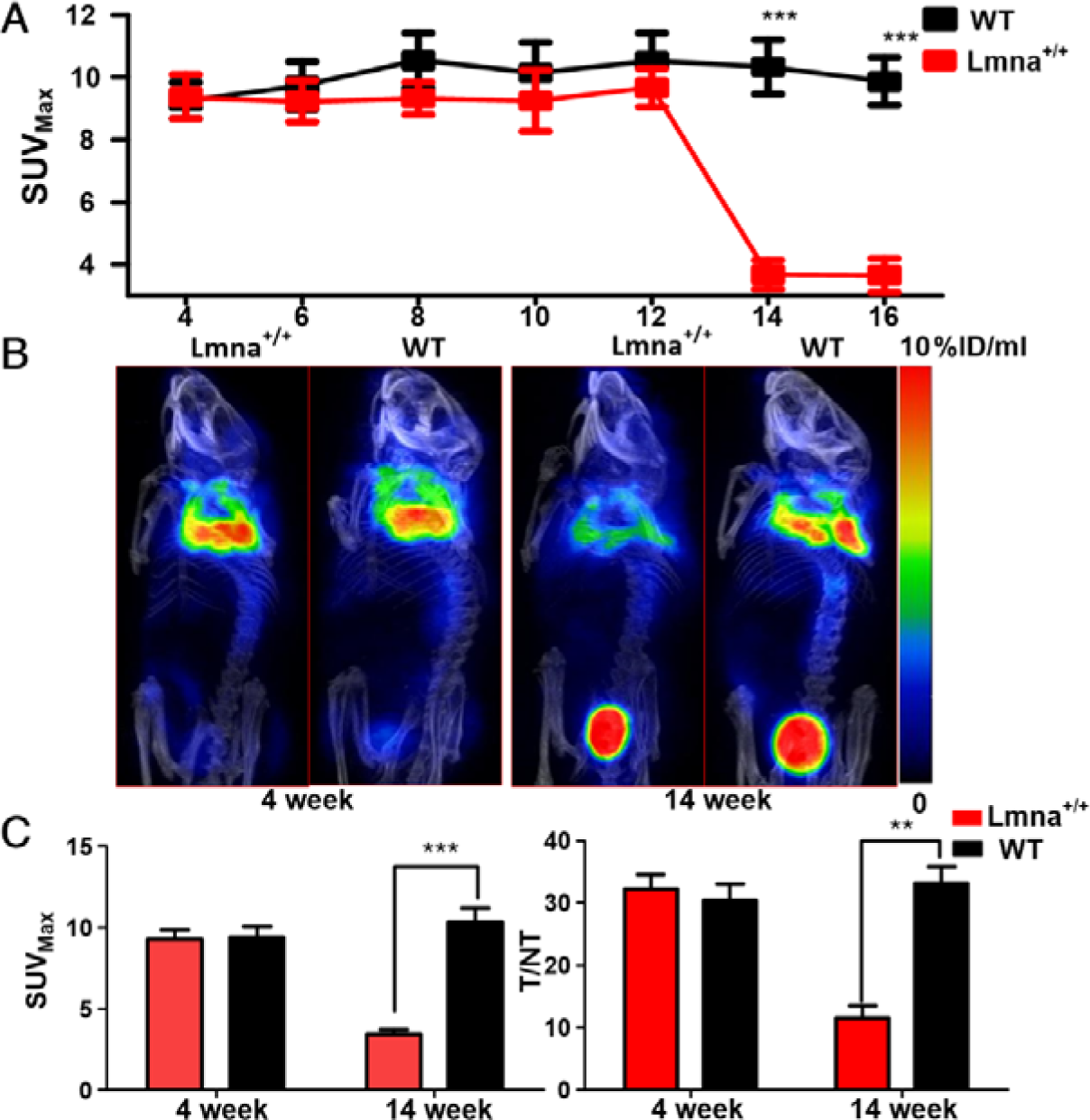
PET of BAT from Lmna^G609G/G609G^ (Lmna^+/+^) mice and wild-type(WT) mice. (A) The changes of SUV_Max_ in BAT from 4 to 16 weeks in mice. (B) Lmna^G609G/G609G^ mice and WT mice PET / CT images at 4 weeks and 14 weeks. (C) Quantification of SUV_Max_ and T/NT (target/nontarget) values of BAT of Lmna^G609G/G609G^ mice and WT mice at 4 and 14 weeks. Lung tissue was defined as NT reference. *P<0.05. **P<0.01. **P<0.001.

### Effects of age on BAT in mice

To clarify the causes of BAT ^18^F-FDG uptake decreased in Lmna^G609G/G609G^ mice, the mRNA and expression of UCP1 and beta3-AR were performed. As shown in Figure 2B, UCP1 and beta3-AR translation in Lmna^G609G/G609G^ mice were significantly lower than that in WT mice at 14 weeks of age (UCP1 0.0474±0.0089 vs. 1.000±0.0666 P=0.0001; beta3-AR0.3143±0.0329 vs. 1.000±0.0445 p=0.0002). As shown in figure2C, at 4 weeks of age, there was no difference in the expression of UCP1 and beta3-AR between Lmna^G609G/G609G^ mice and WT mice (UCP1:1.403±0.1213vs.1.342±0.1959, p=0.8026; beta3-AR: 1.200±0.9090 vs.1.170±0.1703 p=0.8223), while at 14 weeks of age, the expression of UCP1 and beta3-AR was decreased (UCP1: 0.4854±0.1012 vs.1.367±0.2706 p=0.0380; beta3-AR: 0.2830±0.0497 vs.1.445±0.1189 p=0.0008). Immunohistochemistry analysis were consistent with western blot results. These findings confirm that Lmna^G609G/G609G^ mice BAT activity decreased, and appeared of aging performance, which was consistent with changes in aging in normal mice (McDonald and Horwitz 1999).

To study the causes of BAT dysfunction, the expression of PRDM16 and was performed, CDKN2A/p16INK4a was performed, which is regarded to be a biomarker of cellular senescence (Hall et al. 2016), and quantify apoptotic cells in BAT was performed by Terminal deoxynucleotidyl transferase dUTP nick end labeling (TUNEL). As shown in figure 3, the transcription and expression of PRMA16 were decreased at the age of 14 weeks in Lmna^G609G/G609G^ mice (1.000±0.1582 ±. 1.850±0.0945 p=0.0099). There is a greater number of p16INK4a-expressing cells in the BAT of Lmna^G609G/G609G^ mice (64.33±2.333% ±. 50.33±2.603 p=0.0161). Likewise, the number of TUNEL positive cells of BAT in Lmna^G609G/G609G^ mice were more than that in WT mice (4.0±0.45% vs. 1.36±0.1202 p=0.0049).

**Figure 2.**
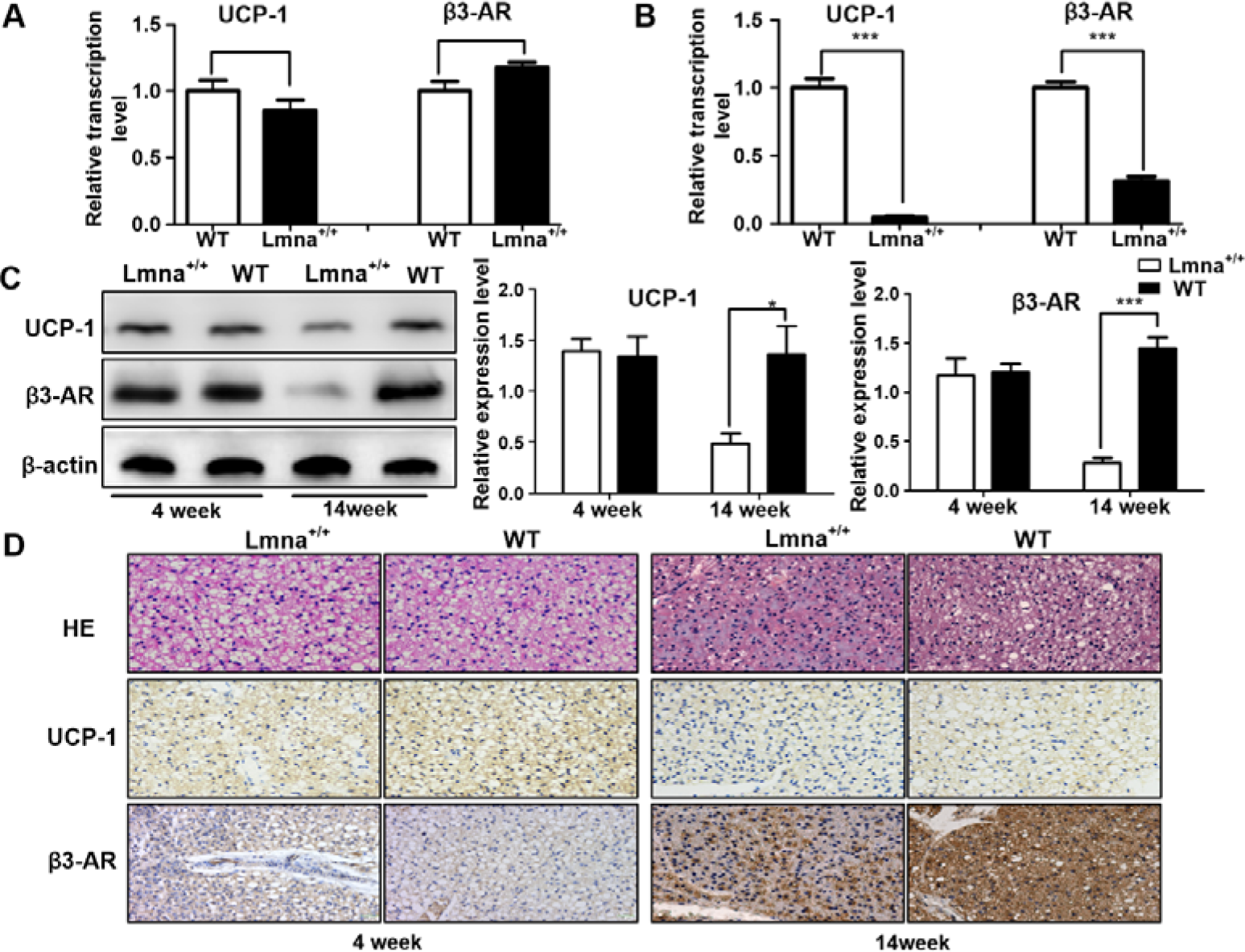
Analysis of beta3-AR and UCP1 level in BAT of Lmna^G609G/G609G^ (Lmna^+/+^) mice and WT mice. (A) (B)Relative transcription level of UCP-1 and β3-AR of aged mice and WT mice at 4week and 14 weeks. (C) Western blot of UCP1, beta3-AR and β-actin and Semi-quantitative analysis of expression levels of UCP1, β3-AR. (D) BAT hematoxylin and eosin stain and Immunohistochemical analysis of beta3-AR and UCP1. *P <0.05. **P <0.01***P<0.001.

**Figure 3.**
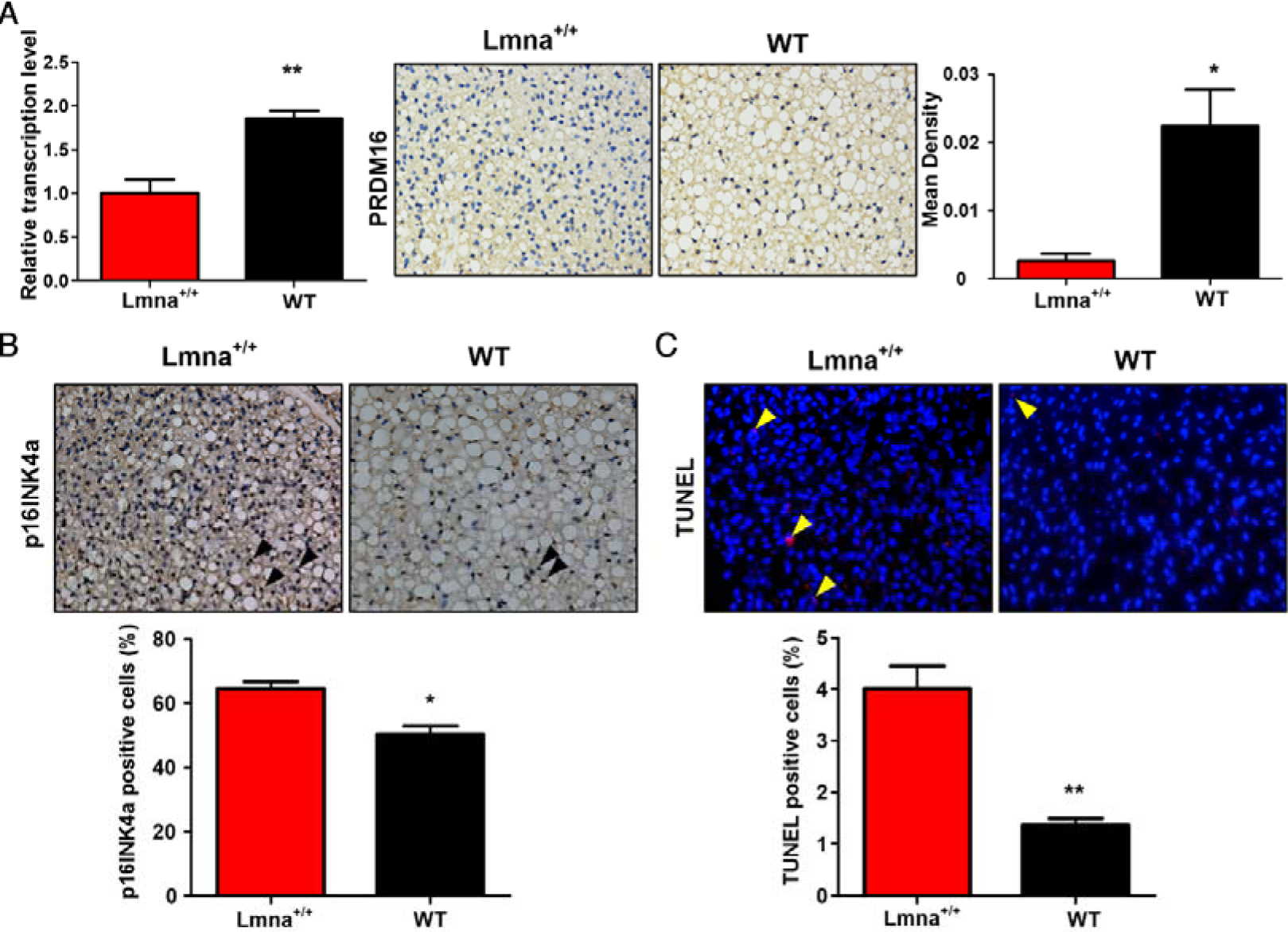
Analysis of PRDM16 level and cell senescence and apoptosis in BAT of Lmna^G609G/G609G^ (Lmna^+/+^) mice and WT mice.(A) Relative transcription level and Immunohistochemical analysis of PRDM16 of Lmna^G609G/G609G^ mice and WT mice at 14 weeks. (B) (C) Images of p16INK4a (black, open arrowheads) immunostaining and TUNEL (yellow, open arrowheads) assay in BAT of Lmna^G609G/G609G^ (Lmna^+/+^) mice and WT mice, and quantification of positive for p16INK4a and TUNEL. *P <0.05. **P <0.01.

### Effects of age on body weight in mice

As shown in figure 4, the body weight of Lmna^G609G/G609G^ mice began to decline in the 10th week, while the weight of wild mice continued to increase, indicating that Lmna^G609G/G609G^ mice began to decline in the state of the body.

**Figure 4.**
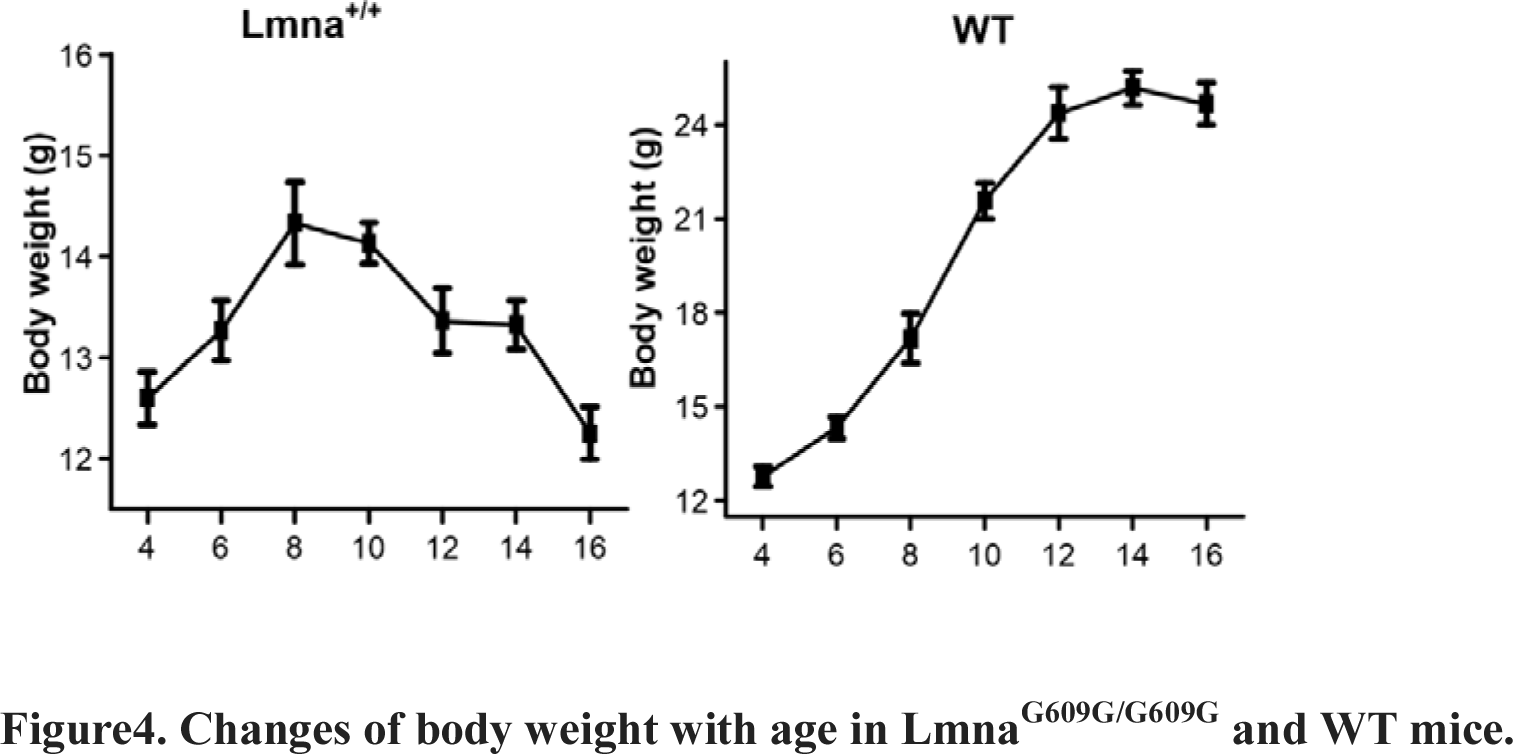
Changes of body weight with age in Lmna^G609G/G609G^ and WT mice.

## Discussion

In this study, we studied the changes of BAT function with age and analyzed the relationship between ^18^F-FDG PET/CT imaging and aging of BAT in Lmna^G609G/G609G^ mice and the mechanism of BAT dysfunction was explored. Our study led to these major findings: Firstly, The BAT function of the Lmna^G609G/G609G^ mice decreased significantly, and the uptake of ^18^F-FDG decreased at the age of 14 weeks. Secondly, Lmna^G609G/G609G^ mice brown adipocytes produce reduced, increased number of senescence and apoptosis. Thirdly, the Lmna^G609G/G609G^ mice began to decline in the state of the body at 10 weeks of age earlier than ^18^F-FDG uptake decrease.

Aging is an irreversible natural process. It is projected that the combined senior and geriatric population will reach 2.1 billion by 2050 (Mba 2010). Aging is the primary risk factor for major human pathologies, including cancer, diabetes, cardiovascular disorders, neurodegenerative diseases (Lopez-Otin et al. 2013), and is one of the most important determinants of BAT activity in all animals, from rodents to humans (Bahler et al. 2016). Therefore, the study of BAT metabolic activity decreased following aging is of great significance. The aging model for the study of BAT is necessary. Whereas, it was not found that the aging model is suitable for BAT studies until now (Hemmeryckx et al. 2012; Vermeij et al. 2016). In our study, we found that Lmna^G609G/G609G^ mice BAT had age-related dysfunction, available for BAT aging research. Lmna^G609G/G609G^ mice for BAT aging study to solve the problem of long study period, inconsistent degree of aging and other issues.

Lmna^G609G/G609G^ mice BAT dysfunction can mimic normal mice BAT aging changes. The changes in BAT in normal mice associated with aging were predominantly reduced in β3-AR and UCP1 expression (Graja and Schulz 2015). We found that the expression of β3-AR and UCP1 in Lmna^G609G/G609G^ mice decreased in a short time and Lmna^G609G/G609G^ mice BAT had age-related dysfunction. The decrease of ^18^F-FDG uptake in BAT of Lmna^G609G/G609G^ mice was closely related to the decrease of β3-AR and UCP1 expression. Exposure to cold has been known to stimulation the sympathetic nervous system via binding of the neurotransmitter norepinephrine to the β3-AR (Himms-Hagen et al. 2000). β3-AR stimulation increases glucose uptake in BAT (Inokuma et al. 2005a) through increasing in transcription and de novo synthesis of glucose transporter molecule 1 (GLUT1) (Olsen et al. 2014). we also found that Lmna^G609G/G609G^ mice GLUT1 transcription level was significantly lower than WT mice (supplementary figure 2). In the cold stimulation, β3-AR not only increases glucose transport, but also increases UCP1 expression and activity (Collins et al. 2010). Although Hankir MK et al. thought that BAT ^18^F-FDG uptake and UCP1-mediated heat production can be dissociated (Hankir et al. 2017), most studies suggest that UCP1 is necessary for norepinephrine-induced glucose utilization in BAT (Inokuma et al. 2005b). The reduction in expression of UCP1 in Lmna^G609G/G609G^ mice further reduced the uptake of ^18^F-FDG in BAT.

The mechanism of BAT dysfunction in Lmna^G609G/G609G^ mice also simulates the mechanism of BAT aging in normal mice (Harms et al. 2014). The mechanism of premature aging in the HGPS mouse model is a point mutation in position 1824 of the LMNA gene, in which cytosine is replaced with thymine. This mutation creates an abnormal splice donor site, which produces a truncated protein (progerin) lacking residues 607-656 of prelamin A but retaining the C-terminal CAAX box, a target for prenylation (De Sandre-Giovannoli et al. 2003b). Progerin cannot be detached from the nuclear membrane, and ultimately damage the structure and function of the nucleus (Cao et al. 2011). Progerin also participated the normal process of aging (Ragnauth et al. 2010; Rodriguez et al. 2009). Therefore, HGPS accelerate a subset of the pathological changes that together drive the normal ageing process (Burtner and Kennedy 2010; Lopez-Otin et al. 2013). Lmna^G609G/G609G^ mice mimic the physiological aging of BAT from both sides. Firstly, Brown fat cells formation was reduced. The expression of PRDM16 expression was decreased in Lmna^G609G/G609G^ mice, which acts as a transcription coregulator that controls the development of brown adipocytes in BAT (Seale et al. 2011). Secondly, the number of BAT cells changed (Sellayah and Sikder 2014). We also found that the increase in BAT cell apoptosis and senescent cells in Lmna^G609G/G609G^ mice resulted in a decrease in the number of cells.

Aging is a systemic change, Lmna mice weight began to decrease in the 10th week. HGPS model were lean (Sullivan et al. 1999), and myopathic disease (Cutler et al. 2002), which lead to body weight decreased before BAT ^18^F-FDG uptake decrease, indicating that Lmna^G609G/G609G^ mice began to decline in the state of the body before BAT aging.

We studied the aging of BAT in Lmna^G609G/G609G^ mice from the perspective of glucose metabolism. ^18^F-FDG is current ‘gold-standard’ for the identification and quantification of BAT metabolic activity, but it cannot discriminate between oxidative and non-oxidative BAT glucose metabolism (Blondin et al. 2015a; Orava et al. 2011). In addition to 18F-FDG, there are still other tracers used in aging of BAT metabolic activity studies (Blondin et al. 2015b; U et al. 2016), to more comprehensive assessment of BAT metabolic activity, the application of other tracers is also necessary.

## Materials and Methods

### Animal Preparation and ^18^F-FDG Micro-PET/CTImaging

Lmna^G609G/G609G^ mice(N=6) obtained from Department of Biochemistry & Molecular Biology, Shenzhen University Health Science Center and experiments were performed in accordance with Animal Experimental Ethics Committee of The Fourth Military Medical University approved protocols. Aging mice were identified by PCR (Supplementary Figure 1). Mice were cold treated under fasting conditions for a duration of 4 hours before they received one dose of ^18^F-FDG (Wang et al. 2012). Ensure the mice are fasting but with access to water. 200-300uCi of ^18^F-FDG in 150ul of saline were intraperitoneal injected into each mouse. The mice were remained in the cold for one additional hour post FDG injection, and then scanned with a small animal-dedicated micro-PET/CT system. Once anesthesia is induced, the animals were moved onto mice bed with its head resting within a cone face mask that continuously delivers Isoflurane (2%) at a flow rate of 1.5 L/min. An electric heating pad is placed under the animal to help maintain the body temperature using a heating pad provided with the small-animal PET/CT system. PET/CT data were acquired for 600s for each mouse with continuous anesthesia. All PET/CT images were processed and analyzed using Nucline nanoScan software (Mediso). For semi-quantitative analysis, three-dimensional (3D) regions of interest were carefully drawn and adjusted manually according to CT images over the borders of the BAT on small-animal PET images of each mice. Three-dimensional round regions of interest were delineated on the lung as a nontarget (NT) reference. Tracer uptake by BAT and lung was quantified as standardized uptake values (SUV) using the formula: SUV=tissue activity concentration (Bq/mL)/injected dose (Bq)×body weight (g). The uptake ratio (T/NT) of the mean BAT and mean NT uptake was calculated and compared.

### Quantitative Real Time Reverse Transcription-PCR

After the mice were sacrificed, total RNA was isolated from BAT using an Total RNA Kit (Omega Bio-Tek, Georgia, USA) according to the manufacturer’s guidelines. Single stranded cDNA was synthesized from total RNA with the PrimeScript™RT Master Mix (TaKaRa, Dalian, China). Real time RT-PCR for each target was performed with SYBR®Premix Ex Taq II (TaKaRa). The thermal cycling condition was set as follows: pre-heating (30 seconds at 94°C) and 40 cycles of d±naturation (10 seconds at 94°C), annealing (30 seconds at 60°C) and elongation (20 seconds at 72°C). Each mRNA level was normalized with the internal control β-actin mRNA level. The pairs of primer are listed in Table 1

**Table 1.**
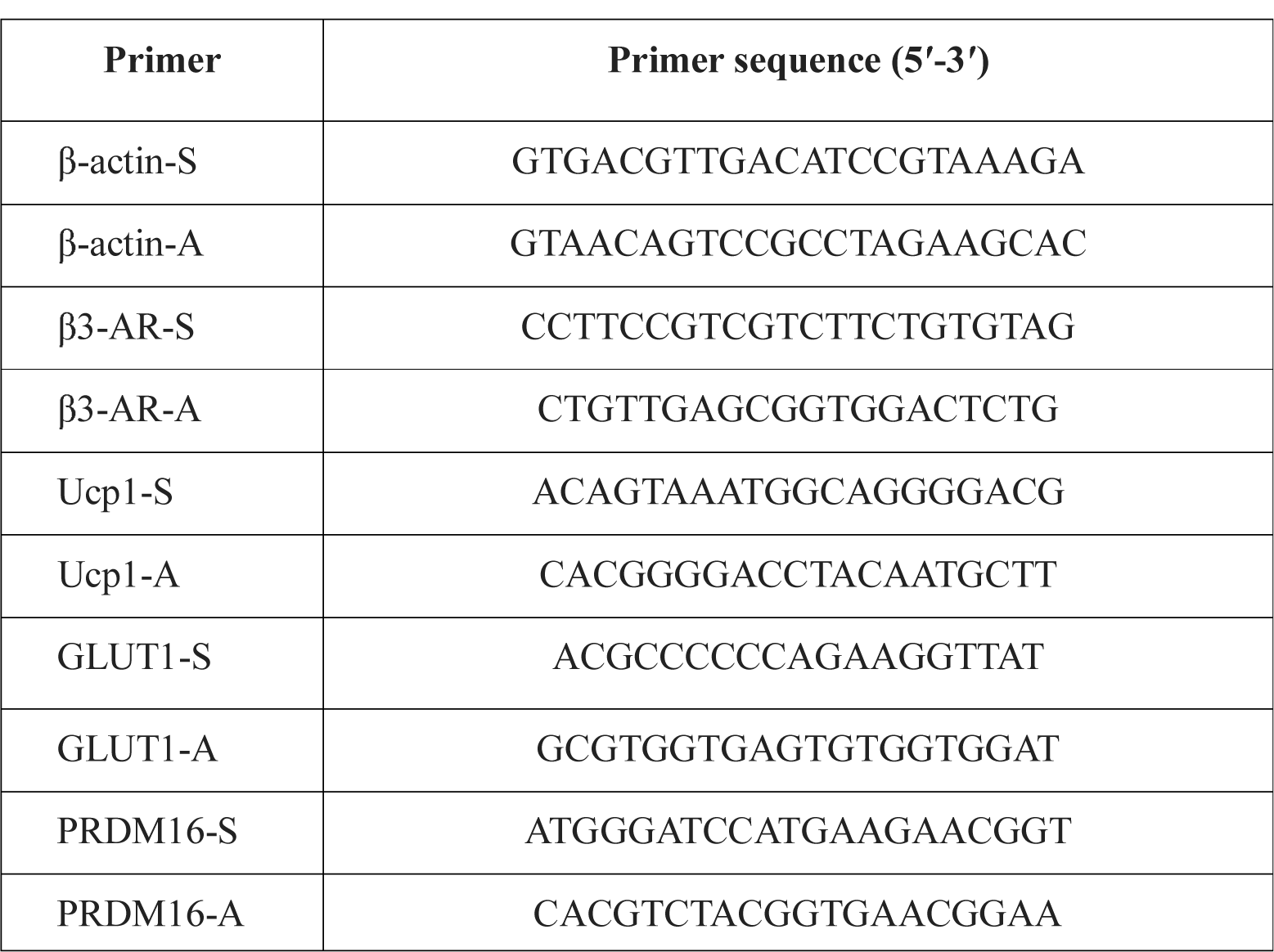
Primers Used in Quantitative RT PCR Experiments

### Western blot analysis

After the mice were sacrificed, protein was extracted, the total protein concentration in the samples was determined using the bicinchoninic acid (BCA; Boster, Hubei, China) method and adding 5×loading buffering heated 5 min at 100. 40μg of protein from each sample was loaded onto 10% sodium dodecyl sulfate-polyacrylamide 1-mm gels and transferred onto NC membranes. To detect the target proteins, we blocked the membranes for 2h with 5% bovine serum albumin (BSA; Boster, Hubei, China) at 4. Then, the membranes were incubated overnight at 4°C with anti-beta3(1:1000, Abcam, USA) antibodies or anti-UCP 1(1:1000, Cell Signaling Technology, USA) specifically recognizing the target proteins. The membranes were subsequently washed with Tris-buffered saline containing Tween-20 (TBST) for 5 min five times and incubated for 1 h at room temperature with secondary antibody. The membrane was washed in TBST for 5 min five times. Blots were detected using enhanced chemiluminescence (ECL) method. The images were captured and analyzed by ImageJ software.

### Immunohistochemistry Analysis

The BAT was harvested from dead mice and fixed in 10% formalin. Formalin-fixed, paraffinembedded tissue blocks were serially cut into 3-mm-thick sections, which were dewaxed in xylene and rehydrated through a graded series of ethanol solutions. After 3 washes in PBS, heat-induced antigen was retrieved in 0.01 M citric acid buffer (pH 6.0) and autoclaved for 5 min at 120. Nonspecific binding sites were blocked through preincubation with normal bovine serum for 30 min. Slices were washed 3 times in PBS for 5 min each wash. These tissue sections were then incubated with anti-beta3-AR antibodies (1:100; Abcam), anti-UCP1 antibodies (1:50; Cell Signaling Technology), PRDM16 (1:100; Abcam), or p16INK4a (1:500; Abcam, diluted in 4% BSA dissolved in PBST 2.5% Triton X-100 in PBS) followed by horseradish peroxidase-conjugated antirabbit IgG (1:1000; EarthOx). In all sections, positive cells were visualized using 3,3-diaminobenzidine tetrahydrochloride (Shanghai Sangon) as a chromogen and were counterstained with hematoxylin. Quantification of the immunostaining was performed by digital image analysis with the Image-Pro Plus 6.0 software (Media Cybernetics). The integrated optical density (IOD) of all the positive staining in each field and areas of interest (AOI) were measured. The IOD was used to evaluate the area and intensity of the positive staining. The mean density (IOD/AOI) represented the concentration of specific protein per unit area.

### Detection of Cell Apoptosis in Brown Adipose Tissue

Tissue apoptosis was assessed using sections for Tunel staining (Beyotime, Beijing, China) according to the manufacturer’s protocol. Briefly, after dewaxing and hydration, sections were incubated with TUNEL reaction mixture. Nuclei were stained using DAPI. Afterwards, slides were observed using a fluorescence microscope (400×; TE-2000U, Nikon, Japan). For each staining, totally 3 sections per group are observed.

### Statistical Analysis

All values are expressed as mean ± SEM. Statistical analysis was performed using GraphPad Prism Software. The differences between two groups were determined by Student’s t test. The probability value of P < 0.05 was considered significant.

## Acknowledgments

We thank Liu Yang from Department of Orthopaedics, Xijing Hospital, Fourth Military Medical University, (Xi’an, China) for her generous support.

## Conflict of interest statement

There is no conflict of interest in this manuscript.

## Funding

This work was supported by the National Natural Science Foundation of China (Grant Nos. 81230033, 81227901), the National Key Research and Development Program of China (Grant No. 2016YFC0103804) and supported by Research Grants from Shenzhen (CXZZ20140903103747568, KQTD20140630100746562 and Discipline Construction Funding of Shenzhen)

## Supplemental material

**Supplementary figure 1.**
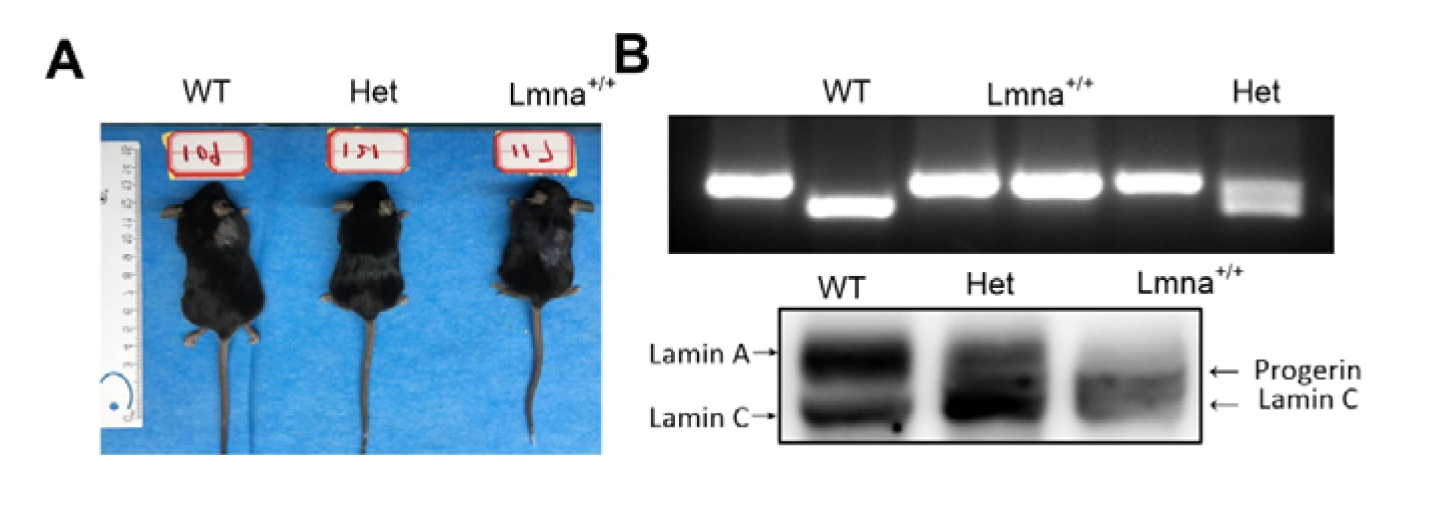
Identification of Lmna^+/+^ mice and characterization of the model. (A) Comparison of mice at 14 weeks of age. (B) Lmna^+/+^ mice were identified by PCR and western blot.

**Supplementary figure 2.**
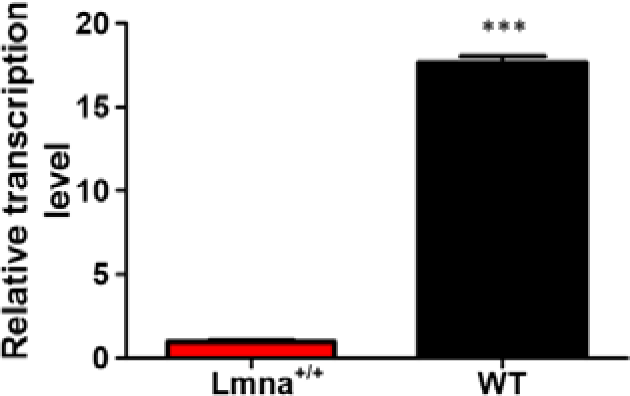
Relative transcription levels of GLUT1 in the BAT of Lmna^+/+^ and WT mice. *** P < 0.001.

